# Glioblastoma Invasion Remodels Neural Circuits and Drives Persistent GABAergic Dysfunction in Human Brain Organoids

**DOI:** 10.64898/2026.08.11.744022

**Authors:** Ewa Grassin, Himanshu Chintalapudi, Xianjun Dong, David S. Goldman, Danielle Hagee, Chunxiao Cui, Aaron Goldman, Luke P. Lee

## Abstract

**Background:** Glioblastoma (GBM) is characterized by neurological dysfunction caused by tumor cells that interact with and alter neuronal circuits. However, the specific neuronal populations and molecular mechanisms most susceptible to GBM invasion remain poorly understood.

**Methods:** We created a human tumor–brain organoid model by combining U87 glioblastoma cells with iPSC-derived neural organoids. This system enabled us to study tumor–neural interactions over an extended period under standard temozolomide (TMZ) treatment. We used single-cell transcriptomics to monitor cell-type-specific responses.

**Results:** Our model recapitulated the diffuse infiltration observed in patients, leading to extensive structural remodeling and a profound loss of neuronal and glial populations. Single-cell analysis revealed that TMZ suppressed proliferative and biosynthetic programs but enriched for stress-responsive, mesenchymal-like, and therapy-adapted tumor states. Notably, GABAergic neurons exhibited the greatest transcriptional vulnerability, with ∼36% (7,499 of 20,659) of genes differentially expressed. Invasion triggered endoplasmic reticulum stress and shut down metabolic, respiratory, synaptic, and ion-homeostatic pathways. Crucially, *SLC12A5*-expressing GABAergic neurons plummeted from 31% to 12%, accompanied by a sharp decline in KCC2 protein expression. While TMZ partially rescued neuronal metabolic and electron transport chain function, it failed to restore *SLC12A5*/KCC2 expression or inhibitory signaling.

**Conclusions:** GBM invasion leads to a continued imbalance of chloride in GABAergic networks, and this disruption remains even after undergoing tumor-targeted chemotherapy. This human iPSC-derived tumor–brain organoid platform provides a reliable and scalable system for studying complex tumor-neural interactions and exploring therapeutic approaches that aim to eliminate the tumor while preserving neural function.

## Introduction

Glioblastoma (GBM) remains the most common and deadliest primary malignant brain tumor in adults, with median survival rarely exceeding 15 months despite aggressive multimodal treatment [1]. Standard-of-care therapy relies on maximal surgical resection followed by radiotherapy and temozolomide-based chemotherapy, and treatment response is influenced in part by MGMT promoter methylation status [2–4]. Despite these interventions, glioblastoma almost invariably recurs, and long-term survival remains uncommon [5]. Moreover, disease progression is frequently accompanied by seizures, cognitive decline, and focal neurological deficits, reflecting the profound disruption of neuronal circuits induced by glioblastoma [6, 7].These persistent challenges underscore the need not only for improved tumor-directed therapies but also for a deeper understanding of how GBM interacts with surrounding brain tissue and contributes to neurological impairment.

A key question in GBM biology is how malignant cells communicate with the normal neural microenvironment. Instead of forming a distinct mass, GBM often infiltrates surrounding neuronal and glial tissues. These two-way interactions are crucial, as they can influence how the tumor grows, invades nearby regions, resists treatments, and leads to neurological issues [8, 9]. Defining these interactions may reveal mechanisms of disease progression that are unique to the brain and identify therapeutic opportunities beyond tumor cell-autonomous vulnerabilities. However, the limited availability of functional human brain tissue has made it challenging to explore these questions through experiments. While animal models remain incredibly useful for understanding tumor growth and treatment effects, they do not fully capture the unique features of the human brain’s structure, cellular composition, and gene activity, especially at the boundary between cancerous and normal tissue [10]. In this context, new approach methodologies (NAMs), including brain organoid systems and other human-based complex in vitro models, provide a powerful platform for investigating tumor–brain interactions in physiologically relevant and experimentally tractable settings [11]. This need is especially meaningful given the growing evidence that neuronal activity is not merely disrupted by GBM but may also contribute to disease progression. Glioma cells can establish connections with neural circuits, respond to neuronal signals, and use synaptic and paracrine communication to promote tumor growth [6, 12, 13]. Clinically, GBM often leads to seizures and other neurological issues related to the tumor, indicating that the malignant tissue impacts the local brain circuits in ways that go beyond just the physical presence of the mass [14] [15]. Although hyperexcitability is a well-recognized feature of glioblastoma-associated neurological disease, the molecular mechanisms by which invasive GBM remodels surrounding human neuronal populations remain incompletely understood[15, 16].

Among the pathways increasingly linked to both neuronal dysfunction and GBM biology is inhibitory neurotransmission mediated by the neuronal K+-Cl− cotransporter KCC2, which is encoded by *SLC12A5*.

KCC2 is a key regulator of intracellular chloride homeostasis and is essential for hyperpolarizing GABAA receptor responses[17–20]. Loss of KCC2 function has been associated with neuronal hyperexcitability, seizure susceptibility, and impaired inhibitory signaling across multiple neurological disorders [21–24]. Emerging evidence indicates that *SLC12A5*/KCC2 expression is reduced in glioma relative to normal brain tissue and is particularly low in high-grade disease, including GBM[25, 26]. However, whether invasive GBM alters KCC2-centered signaling networks in adjacent human neural tissue remains unknown.

Here, we employed a human tumor–brain organoid model to investigate how GBM invasion affects adjacent tissues over time, analyzing both transcriptional changes and the impact of standard treatments on these responses. Our research has demonstrated that glioblastoma multiforme (GBM) cells progressively acquire infiltrative characteristics within the organoid model, resulting in considerable neuronal loss and substantial alterations in gene expression as observed at the single-cell level. These effects were particularly pronounced in GABAergic neurons, where tumor exposure activated the endoplasmic reticulum stress-related pathway and inhibited metabolic, respiratory, ion-homeostatic, and postsynaptic signaling networks. A key part of these changes was the reduction of *SLC12A5*/KCC2 expression, which was confirmed at the protein level. Although temozolomide partially restored specific stress-response mechanisms and electron transport pathways, several challenges persisted, notably in the regulation of ion balance and inhibitory signaling, which continued to pose significant concerns. These findings emphasize how GBM invasion can interfere with inhibitory signaling. They further introduce a human organoid platform to investigate tumor–brain interactions. Additionally, this platform presents new opportunities to develop therapies that preserve the integrity of inhibitory circuits, thereby fostering more promising and accessible treatment modalities.

## Results

### Single-cell profiling of a human glioblastoma–brain organoid model reveals selective vulnerability of neurons to tumor invasion

To establish a humanized model of glioblastoma–brain interactions, we generated brain organoids from human induced pluripotent stem cells (iPSCs) and co-cultured them with U87 human glioblastoma cells (Figure 1A). Organoids were matured for over three months prior to tumor exposure, permitting the development of structured neural tissue with established neuronal and glial populations, as confirmed by histochemical analysis (Supplementary Figure 1A). Mature organoids (day 96 of development) were mixed with U87 human glioblastoma cells, which were applied as single cells but quickly aggregated into spheroids over the course of 24 hours. After 5 days, GBM spheroids were assembled into brain organoids, which developed visible outgrowths that progressively enlarged (Figure 1B). By day 12, immunofluorescence imaging revealed extensive tumor infiltration of the organoid parenchyma, as indicated by proliferating GBM cells (Ki67⁺) relative to MAP2⁺ neuronal regions, suggesting direct interactions between tumor cells and host neural tissue (Figure 1B, C).

**Figure 1.**
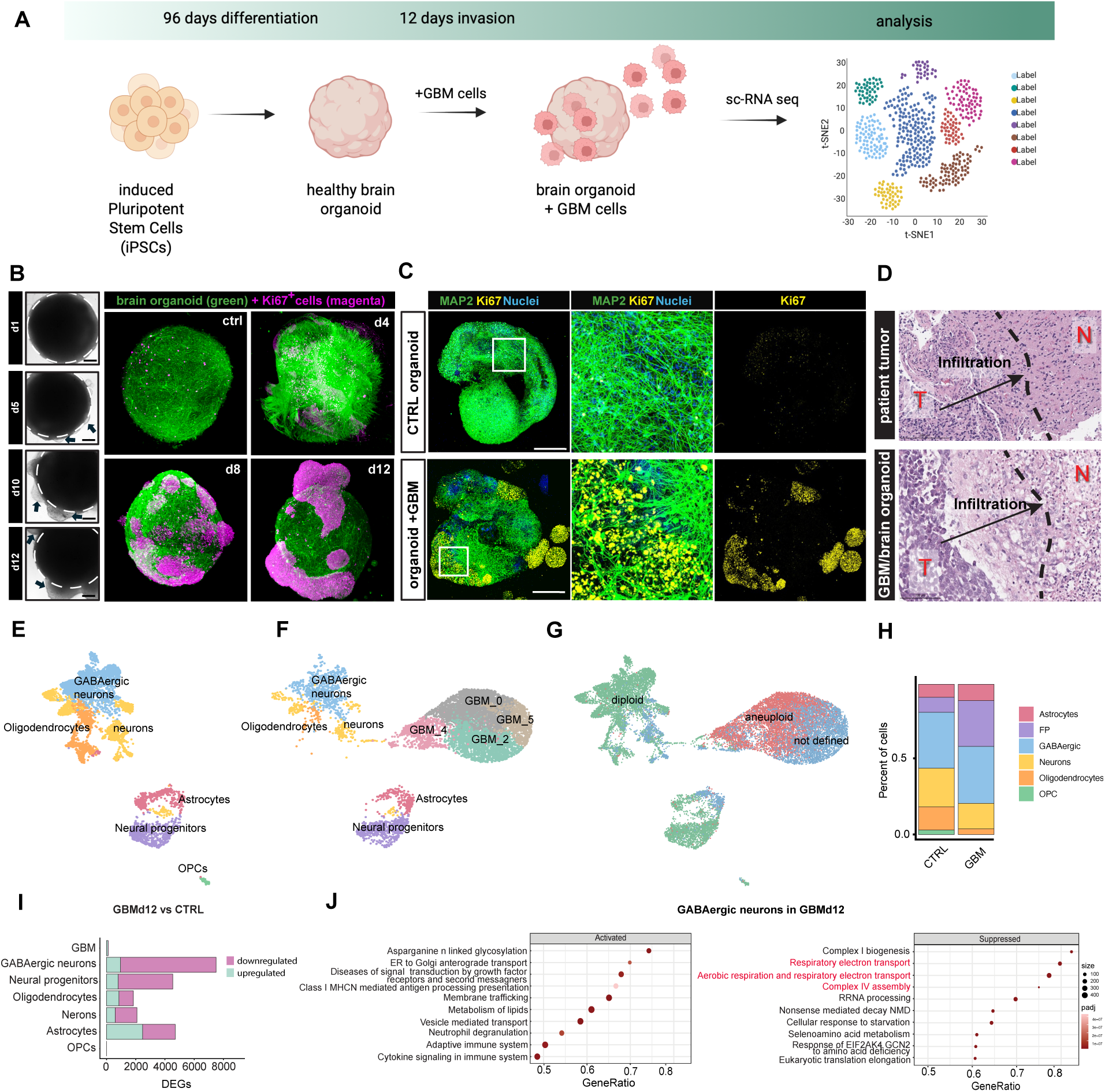
Glioblastoma infiltration into brain organoids reshapes transcriptional programs and disrupts inhibitory neuronal homeostasis. (A) Schematic overview of the experimental workflow. Human iPSC-derived brain organoids at day 96 of development were co-cultured with glioblastoma (GBM) single cells for 12 days. Single-cell RNA sequencing was performed for all three conditions at the end time point. (B) Brightfield images of GBM-invaded organoids on days 1, 5, 10, and 12 post-tumor addition. Progressive tumor invasion is evident, with visible alterations in the organoid surface and increasing integration of tumor masses over time. 3D image of brain organoid (green) with cancer cells (pink). Scale bar 500 µm. (C) Whole-mount immunofluorescence staining of organoids before and after GBM addition. Scale bar 500 µm. MAP2 (green) marks mature neurons and Ki67 (yellow) highlights proliferative cells. Post-GBM addition, increased Ki67 expression and disruptions in MAP2-positive neuronal architecture are observed, indicating tumor-associated proliferation and structural remodeling of the neuronal network. (D) Representative histological images comparing the invasive front of a human glioblastoma specimen (top) and the GBM–brain organoid model (bottom). In both cases, tumor cells (T) infiltrate the adjacent neural tissue (N), recapitulating the diffuse infiltrative growth pattern characteristic of human glioblastoma. The dashed line indicates the approximate tumor–brain interface, and arrows denote the direction of tumor infiltration. (E) Uniform Manifold Approximation and Projection (UMAP) visualization of single-cell transcriptomics from a control brain organoid at day 108 of development. (F) UMAP visualization of single-cell transcriptomics 12-days post GBM-invasion into brain organoid. (G) CopyKAT visualization of integrated single-cell transcriptomic profiles from three samples, revealing diploid populations corresponding to healthy brain cells and aneuploid populations corresponding to GBM cells. (H) Cellular population proportion before and after the addition of GBM cells (I) Differential Gene Expression Analysis of GBM-invaded brain organoid across different cell types: FP – Floor Plate Progenitors, OPC – Oligodendrocyte Progenitor Cell. (J) Gene Ontology (GO) enrichment analysis of GBM in brain organoid (GBMd12 vs CTRL) showing activated and suppressed pathways. (K) GO enrichment analysis revealed significant dysregulation of ion homeostasis-associated pathways. (L) *SLC12A5*, gene encoding KCC2, expression in GABAergic neurons before and after GBM invasion into the brain organoid. (M) KCC2 expression at the protein level N) Quantification of KCC2+ cells in control and GBM-invaded brain organoids across 5 sections from 3 biological replicates.

Using hematoxylin and eosin (H&E) staining, we compared patterns of tumor infiltration in organoids with patient tumors, and analysis by a clinical pathologist confirmed similar histologic features of invasion (Figure 1D). These observations suggested that the tumor-neural organoid model can faithfully reproduce key histopathological features of glioblastoma invasion, providing a physiologically relevant platform for studying the molecular architecture of tumor–brain interactions.

To investigate whether the pathological features were accompanied by changes in cellular composition and transcriptional programs, we performed single-cell RNA sequencing of control brain organoids (Ctrl) and the tumor-brain organoids exposed to GBM cells for 12 days (GBM). Clustering and marker-based annotation identified major neural populations in control organoids, including neural progenitors, mature neurons, GABAergic neurons, astrocytes, and oligodendrocyte lineage cells (Figure 1E). In GBM-exposed organoids, copy number analysis with CopyKAT distinguished aneuploid tumor cells from diploid neural populations, confirming the identity of GBM cells (Figure 1F, G). Notably, GBM invasion led to a cellular reorganization of organoids, primarily by reducing the proportions of neurons, oligodendrocytes, and OPCs relative to those in control brain organoids (Figure 1H).

### GABAergic neurons and the KCC2 axis exhibit a strong transcriptional response to GBM invasion

To understand the molecular and transcriptional impact of GBM on normal neuronal circuits, we interrogated cell-type–resolved differential gene expression. Analysis revealed pronounced transcriptional changes across neural populations following GBM invasion. Notably, among all cell types, GABAergic neurons exhibited the strongest transcriptional response. In total, 20,659 differentially expressed genes (DEGs) were detected at day 12 post-invasion, with a substantial fraction (7,499 DEGs, ∼36%) enriched in GABAergic neurons (Figure 1I). Pathway enrichment analysis revealed widespread disruption of neuronal homeostasis in this population. Genes involved in mitochondrial oxidative phosphorylation and ATP production were significantly downregulated, indicating impaired bioenergetic capacity (Figure 1J). In parallel, stress-response and proteostasis pathways were activated, including ER stress regulators and molecular chaperones. Pathway enrichment analysis revealed many perturbations in neuronal signaling, tissue organization, RNA and protein processing (Supplementary Figure 1B), with inhibitory neurons displaying the strongest transcriptional perturbations, suggesting that they represent a particularly vulnerable neuronal population during tumor invasion.

To identify molecular features linking these coordinated transcriptional changes, we performed gene ontology enrichment analysis of differentially expressed genes in GABAergic neurons, focusing on the biological processes most strongly altered by GBM invasion. Comparison of the genes contributing to these enriched terms identified *SLC12A5*, encoding the neuronal potassium–chloride cotransporter KCC2, as a recurrent component of pathways governing ion homeostasis, osmotic regulation and postsynaptic organization (Figure 2A). This convergence nominated KCC2 as a candidate organizing node within the disrupted inhibitory-neuronal program. Indeed, *SLC12A5*, the gene encoding KCC2, was transcriptionally reduced in GABAergic neurons from GBM-invaded organoids compared with controls, accompanied by a marked decrease in the fraction of expressing cells (31% in control organoids versus 12% following GBM exposure) (Figure 2B). Consistent with these transcriptomic findings, immunofluorescence staining confirmed reduced KCC2 protein levels within neuronal regions of GBM-invaded organoids relative to control organoids (Figure 2C, D).

**Figure 2.**
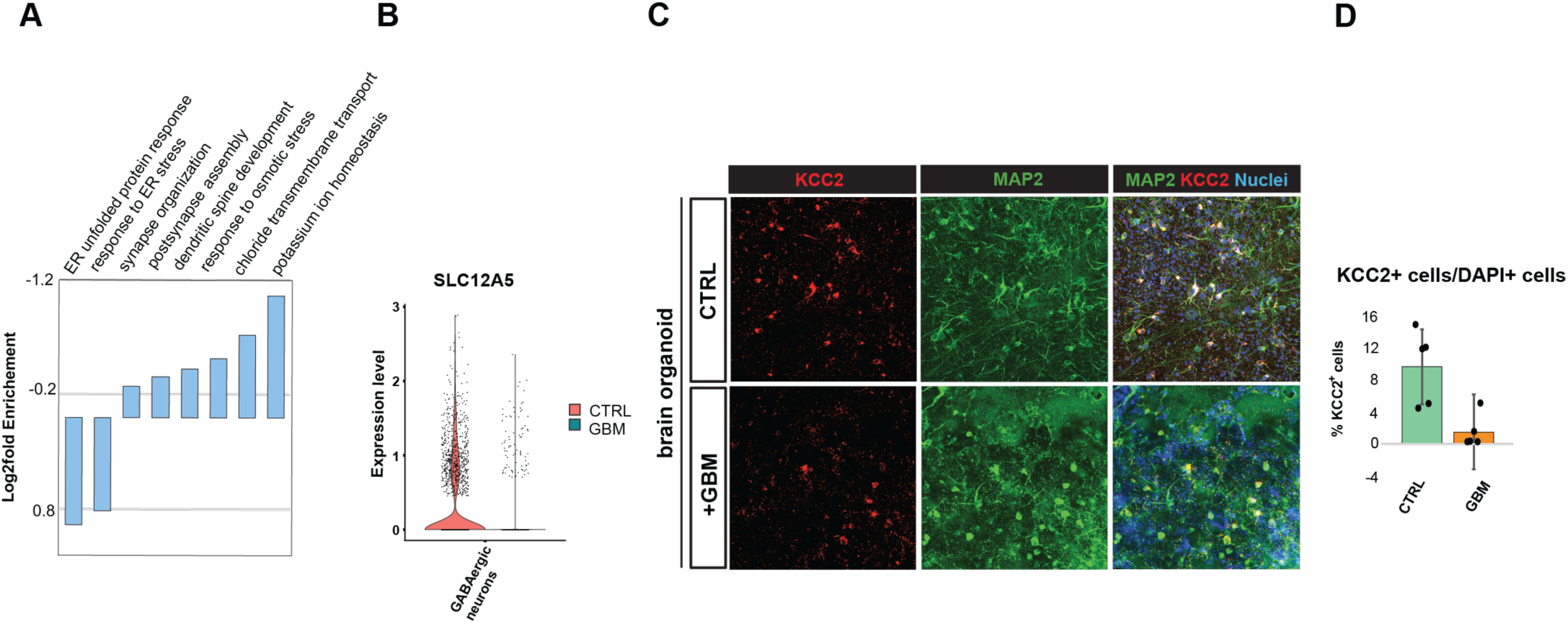
GBM invasion disrupts inhibitory neuronal homeostasis and identifies KCC2 downregulation as a potential molecular mechanism connecting tumor invasion to neuronal dysfunction. (A) GO enrichment analysis revealed significant dysregulation of ion homeostasis-associated pathways. (B) *SLC12A5*, the gene encoding KCC2, expression in GABAergic neurons before and after GBM invasion into brain organoid. (C) KCC2 expression at protein level D) Quantification of KCC2+ cells in control and GBM-invaded brain organoid across 5 sections of 3 biological replicates

Collectively, these findings suggest that GBM invasion disrupts inhibitory neuronal homeostasis and identify KCC2 downregulation as a potential molecular mechanism connecting tumor invasion to neuronal dysfunction. The recognition that GABAergic neurons and the KCC2 axis are highly responsive to GBM invasion shifts the research focus from excitatory neurons to the inhibitory GABAergic system. It also reveals KCC2-mediated defense mechanisms in healthy tissue that are activated during glioblastoma invasion. Human brain organoids enhance our understanding of how tumor invasion affects chloride regulation and help identify a new molecular target for future therapeutic interventions, thereby opening new avenues for ongoing research.

### Temozolomide (TMZ) suppresses proliferation while inducing stress-adaptive GBM cell states within tumor–neural organoids

Temozolomide (TMZ) is a standard-of-care alkylating agent for glioblastoma, with clinical benefit greatest in tumors harboring MGMT promoter methylation [3, 27]. Nevertheless, resistance frequently emerges, and the cell-state transitions accompanying early therapeutic adaptation remain incompletely understood [28]. We therefore asked how TMZ reshapes tumor and neural populations within the tumor–neural organoid model, using U87 cells as a TMZ-sensitive GBM population. After 12 days of tumor invasion, organoids were exposed to TMZ for 30 h and subsequently analyzed by single-cell RNA sequencing (Figure 3A).

**Figure 3.**
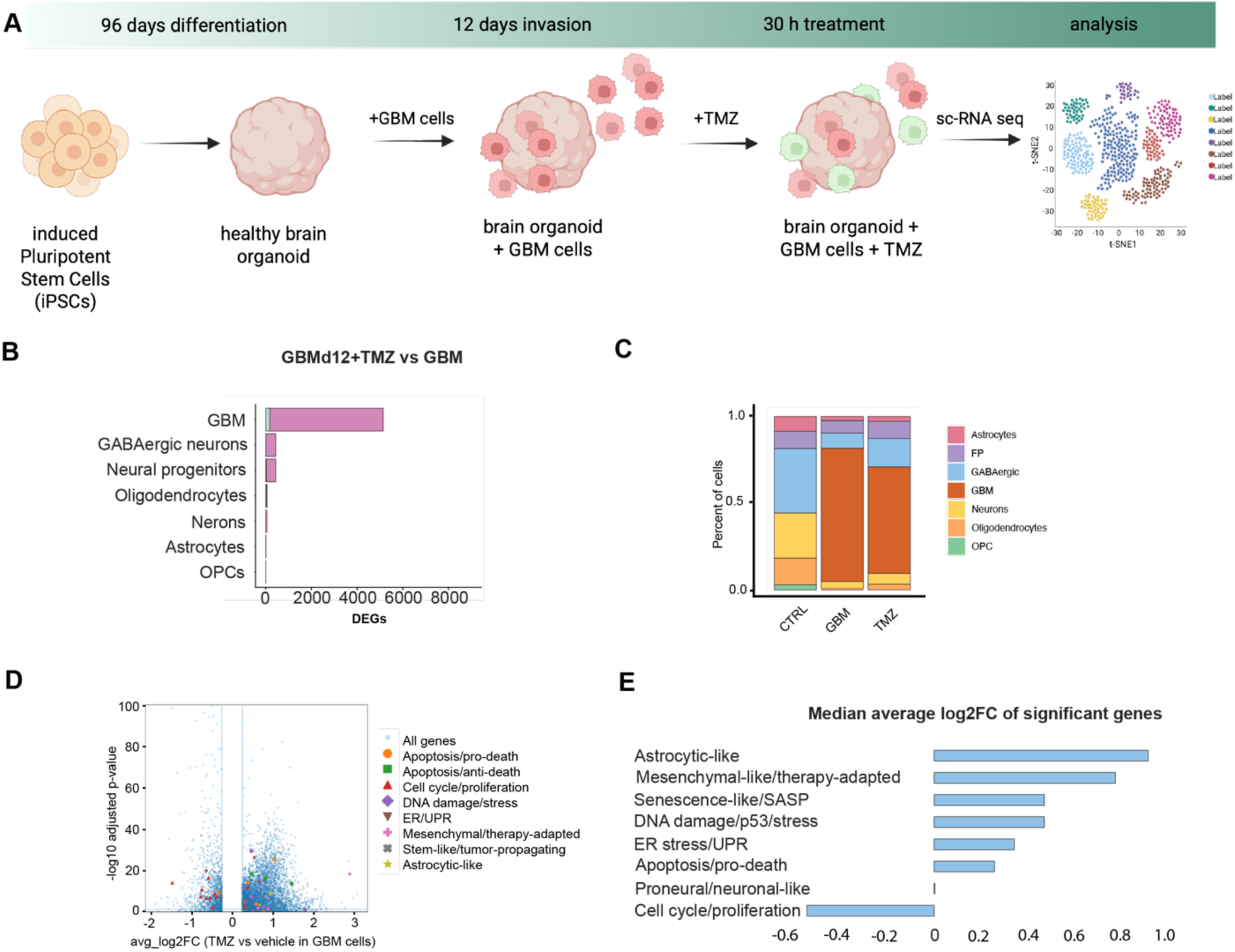
Differential transcriptional and pathway responses to temozolomide in glioblastoma versus healthy cell populations. (A) Schematic representation of experimental setup. (B) Number of differentially expressed genes in GBMd12 + TMZ vs CTRL and GBMd12 + TMZ vs GBM across all cell types. (C) Cellular populations proportions for CTRL, GBMd12, and GBMd12 + TMZ. (D) Volcano plot showing differential gene expression in GBM cells following TMZ treatment compared with GBM. Selected genes associated with apoptosis, cell-cycle regulation, DNA damage/stress responses, endoplasmic reticulum stress/unfolded protein response (ER/UPR), mesenchymal/therapy-adapted states, stem-like/tumor-propagating programs, and astrocytic-like states are highlighted. The x-axis indicates average log2 fold-change (TMZ vs. GBM), and the y-axis indicates –log10 adjusted p-value. (E) Median average log2 fold-change of significantly differentially expressed genes grouped by functional categories and GBM cell-state programs.

As expected, GBM cells exhibited the most extensive transcriptional response to TMZ, characterized by widespread gene downregulation relative to the neural populations (Figure 3B). At the pathway level, TMZ suppressed biosynthetic and proliferative processes, including RNA metabolism, translation and cell-cycle-associated programs (Supplementary Figure 1B). TMZ treatment also reduced the relative representation of GBM cells within the organoids, consistent with acute suppression of tumor-cell expansion in this treatment-sensitive model (Figure 3C).

To define the cell states adopted by the remaining GBM cells, we next examined differentially expressed genes associated with established glioblastoma lineage, stress-response and therapy-adaptation programs (Figure 3D). Consistent with the DNA-damaging activity of TMZ, genes involved in DNA-damage responses, endoplasmic reticulum stress and the unfolded protein response, apoptosis and therapy-associated adaptation were predominantly induced. Astrocytic-like and mesenchymal-like programs were also increased, whereas genes associated with cell-cycle progression and proliferation were broadly suppressed.

We then summarized these responses by calculating the median log2 fold-change among significantly altered genes within each functional category (Figure 3E). Cell-cycle and proliferation-associated programs exhibited the strongest overall reduction following TMZ treatment. By contrast, astrocytic-like, mesenchymal-like and therapy-adapted states, together with DNA-damage, ER stress/UPR and pro-apoptotic programs, showed coordinated upregulation. Thus, acute TMZ exposure reduced GBM abundance and proliferative activity while simultaneously promoting stress-responsive and adaptive transcriptional states among the surviving tumor cells. These findings indicate that, even within a TMZ-sensitive GBM population, treatment rapidly generates heterogeneous cell-state responses that may provide a substrate for subsequent therapeutic resistance.

### Temozolomide (TMZ) partially restores metabolic programs in GABAergic neurons but not KCC2 dysfunction

Because GABAergic neurons exhibited the most pronounced transcriptional perturbations following GBM invasion, we wanted to examine the effects of TMZ treatment within this population as well. In GBM-invaded organoids, GABAergic neurons displayed marked suppression of mitochondrial respiration and electron transport pathways, including Complex I biogenesis and oxidative phosphorylation, consistent with tumor-induced metabolic stress (Supplementary Figure 1C). Following TMZ treatment, pathway enrichment analysis revealed reactivation of aerobic respiration and electron transport programs, accompanied by increased RNA-processing activity, suggesting partial restoration of neuronal metabolic function (Figure 4A).

**Figure 4.**
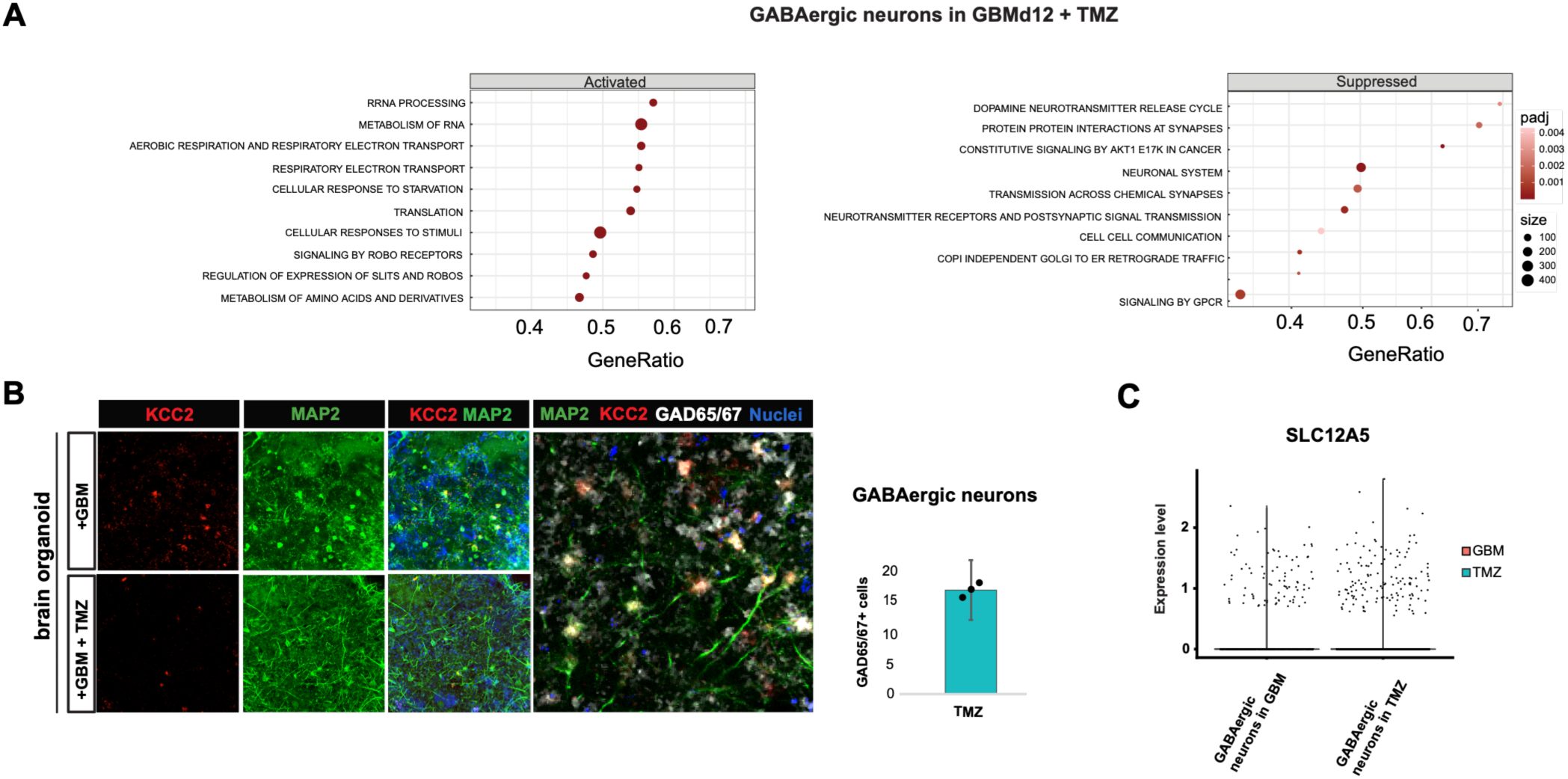
Effect of TMZ treatment on GABAergic neurons. (A) Gene Ontology (GO) enrichment analysis of GABAergic neurons after TMZ treatment. (B) Immunofluorescence staining of GBMd12 organoids before and after TMZ treatment. MAP2 (green) marks mature neurons, and KCC2 (red) highlights the expression of potassium-chloride co-transporter. Immunofluorescence imaging of interneurons (white), KCC2 (red), and general neuronal marker MAP2 (green). Quantification of % of GABAergic neurons relative to nuclei in the TMZ treated sample. (C) *SLC12A5*, gene encoding for KCC2, expression in GABAergic neurons in GBMd12 brain organoid before and after TMZ treatment

Despite these improvements, pathways associated with neuronal signaling, synaptic communication, and KCC2-mediated inhibitory neurotransmission remained suppressed, indicating persistent disruption of ion homeostasis and inhibitory circuit function. Immunofluorescence imaging demonstrated markedly reduced KCC2 protein expression in GBM-invaded organoids, with a further reduction following TMZ treatment. Importantly, GABAergic neurons remained present, indicating that the loss of KCC2 was not simply attributable to the absence of these neurons (Figure 4B). Consistent with these protein-level findings, single-cell RNA sequencing revealed a near-complete loss of *SLC12A5* transcripts in GABAergic neurons relative to control organoids, with no appreciable recovery after TMZ exposure (Figure 4C).

Together, these findings suggest that while TMZ partially restores metabolic programs in GABAergic neurons, it fails to rescue GBM-induced disruption of the KCC2/*SLC12A5* inhibitory signaling axis. These data identify KCC2-centered networks as potential mediators of tumor-associated neural dysfunction and persistent inhibitory circuit impairment.

## DISCUSSION

In this study, we established a human tumor–brain organoid model to examine the reciprocal interactions between invasive GBM cells and surrounding neural populations during tumor progression and under therapeutic pressure. The model reproduced progressive tumor infiltration and revealed broad remodeling of the neural compartment, including loss of neuronal and glial populations and pronounced transcriptional disruption among GABAergic neurons. Acute TMZ treatment suppressed proliferative and biosynthetic programs in GBM, but simultaneously induced stress-responsive, mesenchymal-like, and therapy-adapted states among the remaining tumor cells. In the neural compartment, TMZ partially restored selected metabolic and respiratory programs but failed to reverse disruption of ion homeostasis and inhibitory signaling. Together, these findings indicate that therapeutic response within GBM cannot be understood solely through changes in tumor-cell viability. Rather, treatment reshapes both malignant and non-malignant cell states while leaving persistent abnormalities within the surrounding neural tissue.

These observations also illustrate the increasing importance of new approach methodologies (NAMs) in basic, translational, and preclinical research. Human organoids, microphysiological systems and other complex *in vitro* models can capture tissue-specific cellular interactions that are difficult to resolve using conventional monolayer cultures and may not be faithfully reproduced in animal models because of species-specific differences in cellular composition, development and molecular signaling [29–32]. This is particularly relevant in the brain, where access to viable human tissue is limited and where disease phenotypes emerge from interactions among diverse neuronal, glial and malignant populations. NAMs are therefore positioned not simply as alternatives to animal experimentation, but as complementary experimental systems that can improve mechanistic resolution, support human-relevant biomarker discovery and provide earlier insight into therapeutic efficacy and toxicity [33, 34]. Their growing incorporation into drug-development and regulatory frameworks further emphasizes the need for well-characterized, reproducible and context-specific human model systems [35, 36].

The present model demonstrates one such application and provides the first data set incorporating single-cell-level analysis of malignant and non-malignant treatment responses evaluated simultaneously. These findings should nevertheless be interpreted as an initial mechanistic framework rather than a definitive representation of GBM biology. The current study uses a limited number of experimental conditions and a single established, TMZ-sensitive GBM cell line. U87 cells provide a tractable system for defining proof-of-concept interactions, but they do not capture the genetic, epigenetic and phenotypic heterogeneity of patient tumors [37, 38]. Similarly, iPSC-derived neural organoids vary in cellular composition, regional identity and maturation state, and cannot yet reproduce the complete vascular, immune and structural complexity of the adult human brain [39, 40]. The single-cell findings will require confirmation across additional biological replicates, independent iPSC backgrounds, patient-derived GBM models and molecular subtypes, ideally with integrated statistical approaches that distinguish biological variability from organoid-to-organoid and technical variability. More rigorous bioinformatic analyses, including pseudo-bulk differential expression, gene-set scoring, regulon inference, ligand–receptor analysis and trajectory-based approaches, will also be needed to establish the robustness and directionality of the identified programs. Nevertheless, the convergence of transcriptional and protein-level observations provides important preliminary clues regarding both tumor-cell adaptation and the selective vulnerability of normal neural populations.

Among the neural populations examined, GABAergic neurons exhibited particularly extensive transcriptional perturbation following GBM invasion. Although this does not establish that inhibitory neurons are uniquely vulnerable—as the magnitude of the response may also reflect organoid composition, maturation state or transcriptional detectability—the coordinated suppression of metabolic, respiratory, ion-homeostatic and postsynaptic programs suggests that GBM imposes a substantial functional burden on this population. GABAergic neurons are essential for maintaining excitation–inhibition balance, and their dysfunction may contribute to the hyperexcitability, cognitive impairment and seizure susceptibility associated with infiltrative gliomas [41–43]. Within these altered programs, *SLC12A5* emerged repeatedly across gene sets related to chloride transport, potassium homeostasis, osmotic regulation and postsynaptic organization, with a corresponding reduction in KCC2 protein. KCC2 is a neuron-enriched potassium–chloride cotransporter that maintains the low intracellular chloride concentration required for hyperpolarizing GABA_A receptor signaling in mature neurons [19]. Loss of KCC2 weakens GABAergic inhibition and, under conditions of impaired chloride homeostasis, can render GABAergic transmission depolarizing rather than inhibitory. [17, 44]. KCC2 also contributes to dendritic spine organization, synaptic maturation and network stability through functions extending beyond chloride transport [45, 46]. Its dysfunction has been implicated in epilepsy, neurodevelopmental disorders and neural injury, and reduced peritumoral KCC2 expression has been proposed to contribute to glioma-associated hyperexcitability and seizures [47, 48]. The persistent loss of *SLC12A5* and KCC2 after TMZ treatment therefore suggests that tumor suppression alone may leave the surrounding neural environment vulnerable to continued disinhibition and circuit dysfunction.

KCC2 may also link impaired inhibitory signaling to neuronal survival. Experimental depletion or pharmacological inhibition of KCC2 in mature neurons can activate extrinsic apoptotic signaling and induce neuronal loss, indicating a survival-promoting function that is at least partly separable from its effects on membrane excitability[49, 50]. Thus, reduced KCC2 in GBM-invaded organoids may be more than a marker of disrupted GABAergic signaling and could contribute directly to the selective depletion of mature neuronal populations, placing KCC2 at the intersection of chloride homeostasis, circuit integrity and apoptosis. The relationship between KCC2 and apoptosis resistance in malignant cells is less clearly defined. Reduced *SLC12A5* expression has been associated with glioma progression, and emerging evidence suggests that *SLC12A5* may exert tumor-suppressive effects in glioma cells. [25]. However, it remains unknown whether KCC2 loss directly promotes resistance to apoptosis, reflects a dedifferentiated malignant state, or accompanies broader therapy-adaptive transcriptional programs. The persistence of KCC2-centered disruption alongside the induction of stress-responsive and therapy-adapted states after TMZ raises the possibility that chloride-regulatory pathways influence both neural injury and tumor-cell survival, but the present data do not establish this relationship. Direct, compartment-specific manipulation of *SLC12A5* in neuronal and tumor populations will be required to determine whether KCC2 has distinct or opposing effects on apoptosis, neural function, and therapeutic response.

These findings raise the possibility that restoring KCC2 activity could complement tumor-directed therapy by preserving neural function rather than solely increasing tumor-cell killing. Small-molecule KCC2 enhancers and strategies that increase KCC2 surface stability have reduced neuronal hyperexcitability and improved neurological outcomes in experimental models of epilepsy and central nervous system injury [51–55] . In the context of GBM, such agents could potentially be paired with TMZ or other antitumor treatments to restore chloride extrusion, reinforce inhibitory signaling, and protect vulnerable neuronal populations. Additional approaches might include preventing activity-dependent KCC2 internalization or degradation, modulating upstream regulators of KCC2 trafficking and phosphorylation, or rebalancing neuronal chloride through coordinated targeting of KCC2 and NKCC1. Because these pathways may have different functions in malignant and non-malignant cells, future strategies should prioritize neuron-selective KCC2 augmentation and evaluate whether KCC2 restoration alters tumor growth, invasion or sensitivity to treatment.

## CONCLUSION

In conclusion, we developed a human iPSC-derived tumor–brain organoid platform to model longitudinal glioblastoma invasion and performed single-cell RNA-seq, revealing a selective genetic vulnerability in inhibitory GABAergic neurons during tumor progression. We built a model by combining U87 glioblastoma cells with iPSC-derived neural organoids. This system enabled us to study tumor–neural interactions over an extended period under standard temozolomide (TMZ) treatment. Mechanistically, glioblastoma invasion induces profound depletion of the potassium-chloride cotransporter KCC2, permanently disrupting neuronal chloride homeostasis and circuit signaling. Crucially, we demonstrate a major therapeutic limitation: standard temozolomide chemotherapy fails to rescue this KCC2-driven circuit damage, leaving neural networks unrepaired despite tumor control. Ultimately, this human organoid platform provides a robust translational framework for screening novel, dual-action neuroprotective and anti-tumor therapies capable of restoring inhibitory neuronal function. Although these results are still preliminary, they establish a foundation for mechanistic investigations of tumor–neural interactions and support the use of human organoid-based NAMs to identify therapeutic strategies that effectively manage tumors while preserving neural function.

## MATERIALS AND METHODS

### Generation of human brain organoids

Human induced pluripotent stem cells (iPSCs), ND50086, were obtained from iPSC Neuro Hub at Brigham and Women’s Hospital (BWH)/Harvard Medical School. iPSCs were maintained in mTeSR media (Stem Cell Technologies) on a Matrigel-coated (Corning) tissue culture plate. iPSC colonies were enzymatically dissociated to single cells, and 10,000 cells were plated in each well of an ultra-low attachment 96-well plate with a U-shaped bottom to facilitate homogeneous embryoid body (EB) formation on Day 0. The wells contained neuronal induction medium containing DMEM/F12:Neurobasal (1:1), 1:100 N2 supplement (Invitrogen), 1:50 B27 without vitamin A (Invitrogen), 1% GlutaMAX (Invitrogen), 1% minimum essential media-nonessential amino acid (Invitrogen), and 0.1% β-mercaptoethanol (Invitrogen) supplemented with 1μg/ml heparin (Sigma-Aldrich), 10μM SB431542 (Startech), 1.18 mM LDN193189 (Sigma), 3μM CHIR99021 (BioTechne) and 10μM ROCK inhibitor Y27632 (Stratech). The ROCK inhibitor was added for the first 48 h, and the neuronal induction medium was changed on day 2. As midbrain dopamine neurons originate from the floor plate, on day 4, midbrain organoids were cultured with the addition of SHH C25 II and fibroblast growth factor 8 (FGF8) to initiate the dorsoventral axis. These two morphogens establish an epigenetic Cartesian grid that defines positional information necessary for mDA induction. Around day 7, organoids started to extrude neuroectodermal buds, and at this point, the media was completely removed, and 30 μl of reduced growth factor Matrigel was immediately added to each well using a pipettor equipped with a pre-chilled 200 μl pipette tip. The Matrigel-embedded midbrain organoid was placed into a 37°C incubator for 30 min to allow the Matrigel to solidify and was grown in tissue growth induction medium containing Neurobasal medium, 1:100 N2 supplement (Invitrogen), 1:50 B27 without vitamin A (Invitrogen), 1% GlutaMAX (Invitrogen), 1% minimum essential media-nonessential amino acid (Invitrogen), and 0.1% β-mercaptoethanol (Invitrogen) supplemented with 2.5 μg/ml insulin (Sigma-Aldrich), 200 ng/ml laminin (Sigma-Aldrich), 100 ng/ml SHH-C25II (R&D Systems), and 100 ng/ml FGF8 (R&D Systems) for 24 h. Once the hMLOs were embedded in Matrigel, a more expanded neuroepithelium began to form in the organoids. To promote growth and differentiation, the hMLOs were transferred into ultra-low-attachment 6- well-plates (Costar) by pipetting using a cut 1000 μl pipette tip. The plates contained the final differentiation media, which consisted of Neurobasal medium, 1:100 N2 supplement (Invitrogen), 1:50 B27 without vitamin A (Invitrogen), 1% GlutaMAX (Invitrogen), 1% minimum essential media-nonessential amino acid (Invitrogen), 0.1% β-mercaptoethanol (Invitrogen), 10ng/ml BDNF (Peprotech), 10 ng/ml GDNF (Peprotech), 100 μM ascorbic acid (Sigma-Aldrich), and 125 μM db-cAMP (Sigma-Aldrich). The medium was replaced every 2 days. All cell lines were tested regularly for mycoplasma and were found to be negative.

### Glioblastoma cell culture and co-culture with brain organoids

Glioblastoma U87 cells were obtained from the American Type Culture Collection (ATCC). U87 cells were maintained at 37 °C and 5% CO₂ in Dulbecco’s Modified Eagle Medium (DMEM, Gibco) supplemented with 10% FBS. Cells were dissociated into single cells using Accutase, and 5,000 cells were added to each brain organoid. Co-cultures were maintained for 12 days, at which point all samples were collected for scRNA-seq. In the TMZ condition, GBM- brain organoid co-cultures received 30uM temozolomide treatment for 30 hours immediately prior to the day-12 endpoint.

### Hematoxylin and Eosin (H&E) Staining Protocol

All staining procedures were performed at HistoWiz, Inc, using the Leica Bond RX automated stainer (Leica Microsystems), a Standard Operating Procedure, and a fully automated workflow. Samples are processed, embedded in paraffin, and sectioned into 4μm-thick slices. Formalin-fixed, paraffin-embedded (FFPE) tissue sections were stained using a standardized Hematoxylin and Eosin (H&E) protocol, dehydrated, and mounted using a TissueTek Prisma and Coverslipper (Sakura). Whole-slide scanning was performed at 40× on an Aperio AT2 (Leica Biosystems).

### Whole-mount Immunohistochemistry

The midbrain organoids were fixed in 4% paraformaldehyde (PFA) for 1h. Primary and secondary antibodies were prepared in blocking and permeabilizing solution: 6% BSA, 0.5% Triton-X 100 (Roth), with 0.1% (w/v) sodium azide (Sigma-Aldrich) in PBS (Sigma-Aldrich), and were added for 3 days at 4°C. Between primary and secondary antibody incubation as well as after the staining procedure sample was washed 5 times for 1 hour each with 0.1% Triton X-100 in PBS. This long staining procedure allows the antibodies to fully penetrate the organoid despite their large size and high density. Images were taken on Leica Stellaris confocal microscope.

For the 3D imaging, organoids were prepared with the SHIELD kit according to the manufacturer’s instructions. In short, organoids were fixed in 4% PFA for 1h. SHIELD Epoxy was mixed with SHIELD ON buffer in ratio 1:7. Organoids were incubated in the mixture overnight at 4°C. The samples were then moved to room temperature and incubated for 6h without changing the solution. In the next step, we proceeded with delipidation. Organoids were incubated in Delipidation Buffer at 37°C for 24h followed by washing with PBSTN at room temperature for 3h. Primary antibodies were prepared in PBSTN solution and incubated with organoids for 3 days. Samples were washed for 6 h in PBSTN at 37°C with 5 refreshes of the solution, followed by 2 h 4% fix. Organoids were washed for several hours to remove PFA. Secondary antibody was added in PBSTN solution and organoids were incubated for another 3 days followed by 2h fixation and washes. 3D imaging was performed on Light Sheet Microscope (Zeiss).

### Single cell isolation and RNA sequencing

Five organoids were pooled together and washed with PBS. 5ml Papain was gently mixed with 500 μl DNase, and half of the solution was immediately added to the pooled organoids for a 30-minute incubation at 37°C, 5% CO_2_. After incubation, organoids were broken into smaller pieces by pipetting up and down 30x with a 1 ml pre-cut pipette and placed back in the incubator for another 20 minutes. After incubation, remaining organoids were further broken down into smaller pieces by pipetting 30x with a pre-cut 1ml pipette. The cells were passed through a 40 um cell strainer, spun down at 300 g for 5min, resuspended in resuspension buffer DPBS (Gibco), BSA (Thermo) 1:100, 0.2U/μl RNAse inhibitor (Clontech), and the cell concentration was calculated with AOPI dye using Nexcelom Cellometer K2.

16,000 viable single cells per sample were loaded onto the Chromium controller (10X Genomics) for single-cell partitioning and barcoding. Single-cell cDNA synthesis and amplification of the libraries were performed according to the manufacturer’s protocol (Chromium Next GEM Single Cell 3**ʹ** Reagent Kits v3.1). The molar concentration of libraries was determined with Kapa Library Quantification qPCR (Roche), and size distribution was evaluated using an Agilent Bioanalyzer. Libraries were sequenced on Illumina Novaseq.

### Single-cell RNA-seq data analysis

Cellranger (8.0.1) *count* was run on all FASTQ files to perform read alignment and UMI counting using default parameters. Ambient RNA correction was done using *Cellbender* (0.3.0). All downstream analysis was done with *Seurat* (5.5.0) in R (4.4.3). Cell QC and filtering were done using metrics like number of features, number of UMIs & percentage of reads mapping to the mitochondrial genome (< 10%). Doublet detection and removal was done using the R package *DoubletFinder* (2.0.4). All three samples were integrated using *IntegrateLayers()* in Seurat with the method *Harmony*. After normalization and clustering, the clusters were annotated using the top marker genes generated for each cluster using *FindAllMarkers()* with the “wilcox” method. To infer the tumor clones and differentiate the “normal” cells, copyKAT R package (1.1.0) was used to visualize the predictions of the ploidy for each cell. To infer the cell-cell communication patterns between cell types, the R package *LIANA* (0.1.14) was used with default settings. Differential expression analysis between samples for all cell types was done using *FindMarkers()* function & Pathway analysis on the cell types was done using the R package *clusterProfiler* (1.14.6)

## Declarations

### Ethics approval and consent to participate

Not applicable

## Consent for publication

Not applicable

## Availability of data and materials

The single-cell RNA sequencing datasets generated during this study have been deposited in the Gene Expression Omnibus (GEO) under accession number GSE342886. All other data supporting the findings of this study are available from the corresponding author upon reasonable request.

## Competing interests

A.G has equity interest in Xsphera bio as a consultant

## Funding

This work was supported by the Swiss National Science Foundation through a Postdoc.Mobility Fellowship awarded to E.G. (P500PB_203036).

## Authors’ contributions

E.G. conducted the experiments, analyzed and interpreted the data, prepared the figures, and wrote the manuscript. D.H. and C.C. prepared and processed the clinical samples. D.G. performed the histological analysis. H.C. performed sequencing data quality control and bioinformatic processing and generated scRNA-seq data visualizations. X.D. supervised the sequencing data quality control and bioinformatic analysis. A.G. contributed to data sequencing data visualization, critically revised and edited the manuscript. L.L. supervised the study and critically revised and edited the manuscript. All authors helped to edit and approved the final version of the paper.

## Acknowledgements

We thank Dr Paula Montero Llopis and Dr Adrienne Wells from MicRoN Microscopy Core at Harvard Medical School for providing technical assistance with organoid clearing and 3D imaging. Figure 1A and 3A were created using Biorender.

