## Supplementary Figure 1 for "Glioblastoma Invasion Remodels Neural Circuits and Drives Persistent GABAergic Dysfunction in Human Brain Organoids"

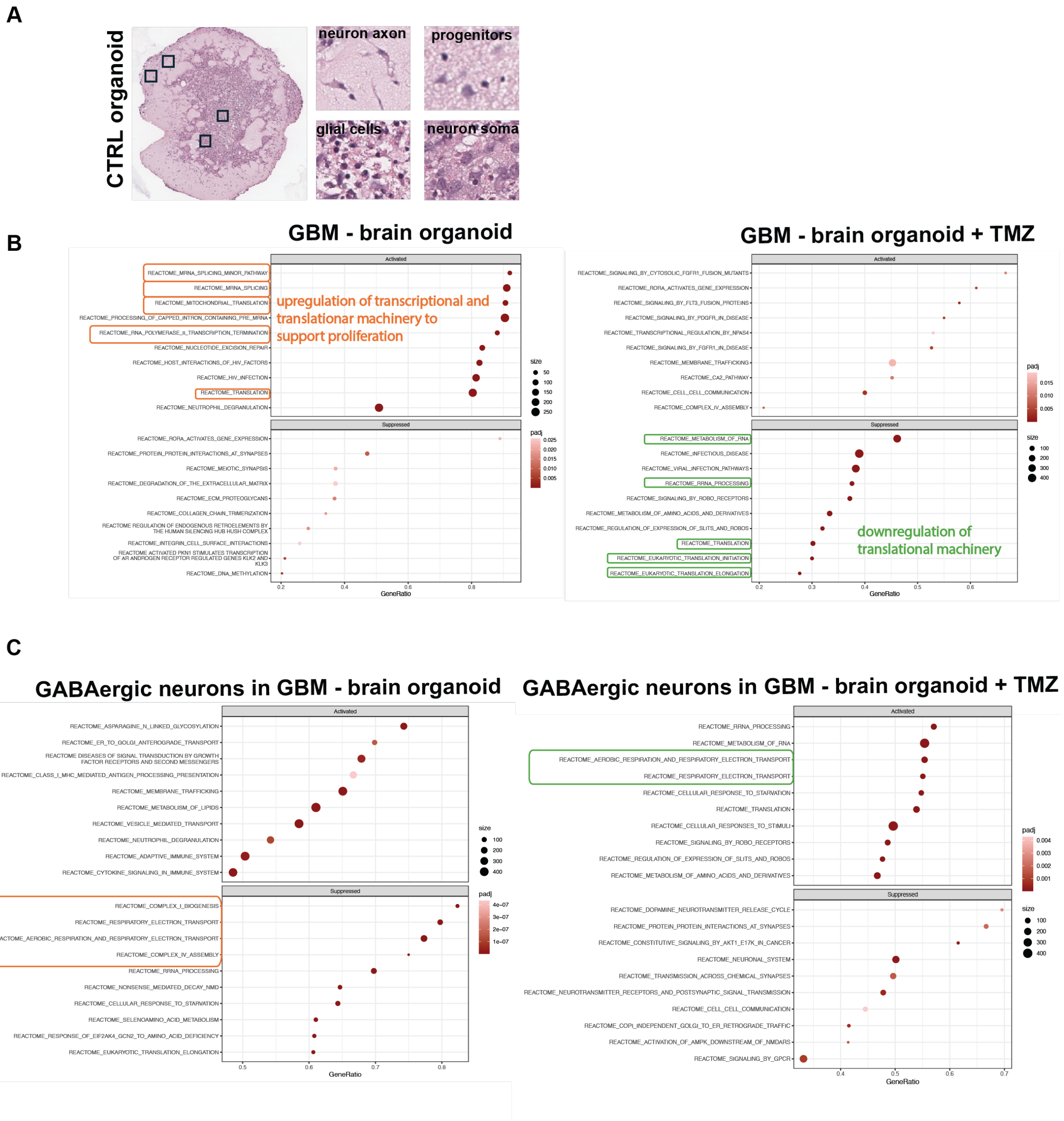


**Figure 1 Histological characterization and pathway enrichment analysis of glioblastoma–brain organoid co-cultures. A)** histological images of control brain organoid. Representative hematoxylin and eosin (H&E)-stained section of a human brain organoid highlighting the major neural structures present in the model. Higher-magnification insets illustrate representative neuronal axons, neuronal soma, glial cells, and neural progenitor cells. **(B)** Reactome pathway enrichment analysis of genes differentially expressed in the whole brain organoid compartment following glioblastoma invasion (left) and after temozolomide

(TMZ) treatment (right). Glioblastoma invasion induced enrichment of pathways associated with transcription, RNA processing, ribosome biogenesis, and translation, consistent with increased biosynthetic activity. TMZ treatment suppressed multiple pathways involved in protein translation and ribosomal function, indicating reduced translational activity following chemotherapy**.** **(C)** Reactome pathway enrichment analysis of differentially expressed genes in GABAergic neurons following glioblastoma invasion (left) and TMZ treatment (right). Glioblastoma invasion suppressed pathways associated with mitochondrial respiration, oxidative phosphorylation, ATP synthesis, and mitochondrial translation, indicating impaired neuronal energy metabolism. TMZ treatment was associated with enrichment of mitochondrial respiratory pathways, suggesting a partial recovery of cellular bioenergetics, while additional neuronal pathways remained altered relative to control.
